# Quantifying Homology and Length Bias in Antimicrobial Peptide Prediction

**DOI:** 10.64898/2026.11.18.767483

**Authors:** Bikash Kumar Paul, Sajib Acharjee Dip, Amarda Shehu, Mohammad Ali Moni, Liqing Zhang

## Abstract

Bacteria are becoming resistant to existing antibiotics faster than new ones are found. Antimicrobial peptides are a promising alternative, but laboratory testing is slow and costly. Candidates are therefore ranked first by a classifier, and published classifiers report near-perfect accuracy. Two controls are missing from the benchmarks behind those numbers. Training and test peptides are not separated by sequence homology. The negative class is not matched to the positive class by length. We propose a controlled evaluation protocol that imposes both. A sample-size control separates matching from the data it removes. We apply it to a four-model ensemble of our own and ten baselines on a benchmark of 16,168 sequences, holding the models fixed. A classifier given nothing but peptide length separates the benchmark far above chance. Homology control costs 0.067 MCC and length matching a further 0.115, taking the reported score from 0.834 to 0.661. Over ten redrawn partitions the length term holds while the homology term shrinks, so length is the effect that survives. The protocol also reorders the methods, and a frozen ESM-2 embedding overtakes the ensemble. Three published benchmarks show the same pattern. We recommend a minimum reporting standard that makes such losses visible.

## I. Introduction

**A**ntimicrobial peptides (AMPs) are short bioactive sequences, typically 5–50 residues, that form part of innate immune defence across most living organisms. Their broad-spectrum activity and comparatively slow induction of resistance make them attractive starting points for therapeutics. Because wet-lab screening is expensive, candidates are first triaged computationally. The resulting literature is large and its reported performance is high. Methods published in the last five years routinely report AUC-ROC above 0.95 on binary AMP and non-AMP discrimination [1]. The three benchmarks we audit in Section IV-K were published with MCC between 0.80 and 0.85, and the ensemble examined here reached AUC-ROC 0.9749. The headline numbers have compressed into a narrow band near the top of the scale.

When a field’s reported metrics saturate, two explanations compete. Either the task is close to solved, or the benchmarks have stopped discriminating between methods. The second explanation is testable, and testing it requires asking what a model would score if it were denied the shortcuts that the benchmark happens to leave open. Two such shortcuts are specific to this problem, and both are left open by common practice.

The first is sequence homology between training and test data. AMP databases are built by curation and expansion around known families, so near-duplicates of a test peptide are frequently present in training. A model that memorises a family will then appear to generalise. Controlling for this requires clustering the benchmark by sequence identity and splitting on clusters rather than on sequences, which is standard practice in protein structure and function prediction, where homology between a query and its training neighbours is treated as something to control for rather than to exploit [2], and less so here.

The second is peptide length, and it is more insidious because it operates through the construction of the negative class. Positives are drawn from AMP repositories such as DBAASP [3], APD [4] and DRAMP [5], which catalogue short peptides. Negatives are typically drawn from UniProt [6] entries lacking an antimicrobial annotation, which are whole proteins and therefore much longer. The resulting benchmark can be separated to a substantial degree by counting residues. Sidorczuk *et al*. [7] established that negative-data selection materially biases AMP benchmarks. What has not been quantified is how a fully specified predictor’s reported performance decomposes once that bias and homology leakage are removed together. Nor is it known whether the relative ordering of methods survives such controls.

This paper proposes an evaluation protocol rather than a predictor. The instrument under test is Deep-AMP, a four-model ensemble we developed on a curated benchmark of 16,168 sequences, which reaches MCC 0.8338 under the conventional random-split protocol. We treat that ensemble and the ten baselines it was compared against as fixed instruments. The models, the features and the code stay unchanged, and only the evaluation protocol varies, so every difference reported here is attributable to the protocol rather than to modelling. Figure 1 shows the benchmark and the leakage its original protocol leaves open.

**Fig. 1.**
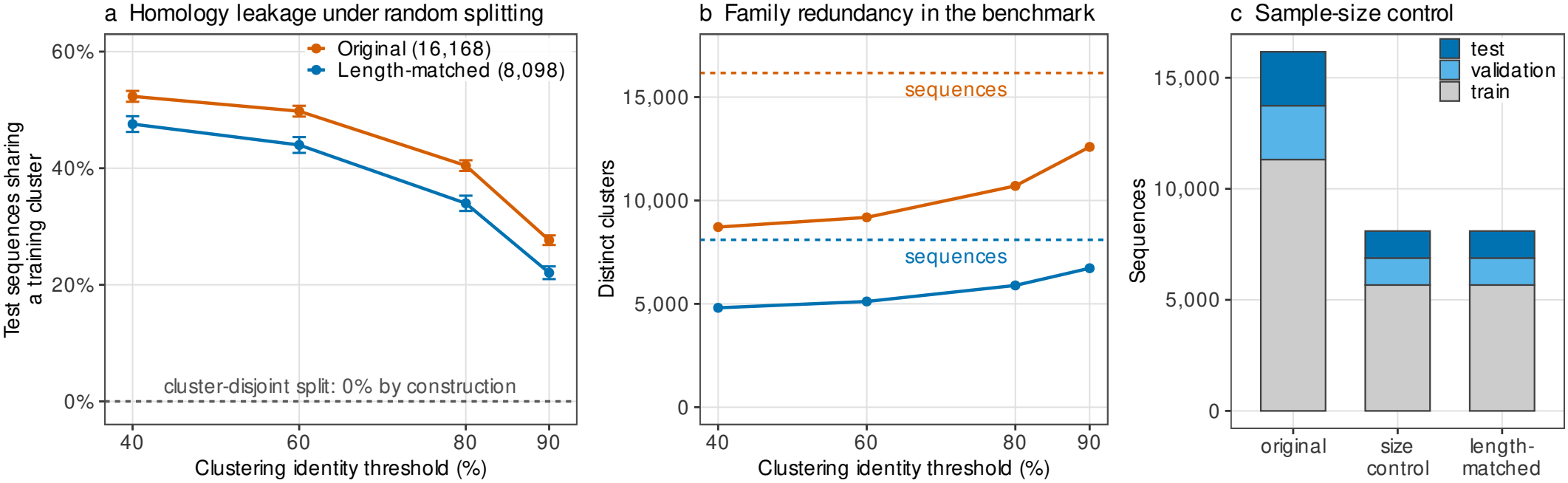
What each control does, measured on the benchmark rather than drawn schematically. (a) Percentage of test sequences sharing an MMseqs2 cluster with the training partition under random 70/15/15 splitting (mean *±* SD over 200 random splits), against the identity threshold at which clusters are defined; the cluster-disjoint partitions used throughout this paper are 0% by construction. (b) Distinct clusters against identity threshold, with each benchmark’s sequence count marked by a dashed line; the gap is the family redundancy that random splitting exposes. (c) Partition sizes for the three benchmarks: the size control matches the length-matched benchmark exactly, separating the effect of length from that of data volume.

The protocol applies three controls and measures each one separately. Cluster-disjoint partitioning at four identity thresholds removes homologous overlap between training and test data. Exact length matching of the negative class removes the length difference between the two classes. A sample-size control separates the cost of matching from the cost of the data it discards. Each configuration is repeated over ten training seeds and over ten redrawn partitions, so that the cost of a control is reported as a distribution rather than as a single number. We then test whether the result is a property of our own benchmark by applying the same controls to three published AMP benchmarks that ship their own splits, and we check the clustering itself against exact Smith-Waterman alignment.

The controls cost the ensemble roughly a fifth of its reported MCC, and that loss is carried mainly by length rather than by homology. They also change which method wins, since the strongest baseline under the original protocol is the one most dependent on the confound, and two models that read sequence directly overtake the descriptor ensemble once the controls are in place. Finally, a classifier given nothing but peptide length recovers roughly half the ensemble’s MCC on the uncontrolled benchmark and falls to chance once lengths are matched. That is the cheapest diagnostic a benchmark can report, and the one we recommend as a minimum.

## II. Background and Related Work

### A. Homology control in sequence-based prediction

Partitioning sequence data by identity-based clustering, rather than at random, is long-established in protein function and structure prediction, where the failure mode, a test sequence whose close homolog was in training, is well documented. Community guidance on supervised learning in biology [8] and on leakage in machine-learning-based science generally [9] both identify identity-aware splitting as a minimum requirement, and reviews of pitfalls in genomic machine learning [10] treat random splits of homologous data as a canonical error. AMP prediction has adopted this practice unevenly; several widely used benchmarks distribute fixed train/test partitions built without identity control, and downstream work inherits them. Concurrent benchmarking work probes the same two controls across released predictors [11], partitioning a single activity corpus by length and by CD-HIT identity at 40%, 60% and 80% to measure each tool’s dependence on them. That study and this one are complementary: it varies the tool and applies each control separately, whereas we hold one instrument fixed and separate the costs of the two controls and of sample size on the same benchmark.

### B. Negative-class construction and the length confound

The AMP task has no natural negative class. Convention is to sample sequences from UniProt that lack an antimicrobial annotation, which introduces two problems. The weaker problem is label noise: un-annotated AMPs will be present among the negatives. The stronger problem, and the one we quantify here, is distributional: AMP repositories catalogue short peptides whereas UniProt entries are predominantly whole proteins, so the classes differ in length by construction. The effect survives the obvious mitigation. The benchmark examined here already restricts its negatives to 50 residues or fewer, and its class medians still differ twofold: 17 against 34 residues (Fig. 2a). The dependence is long-standing: Gabere and Noble [12] found that several web-based AMP classifiers scored long sequences highly largely irrespective of class, and built length-matched negatives in response. Sidorczuk *et al*. [7] later showed that the choice of negative sampling scheme changes measured performance substantially across AMP predictors, and recommended matched negative sets. Our contribution relative to that work is to hold one predictor fixed and measure the full decomposition (homology, length, sample size) on the same instrument, and to test whether method ranking is stable.

**Fig. 2.**
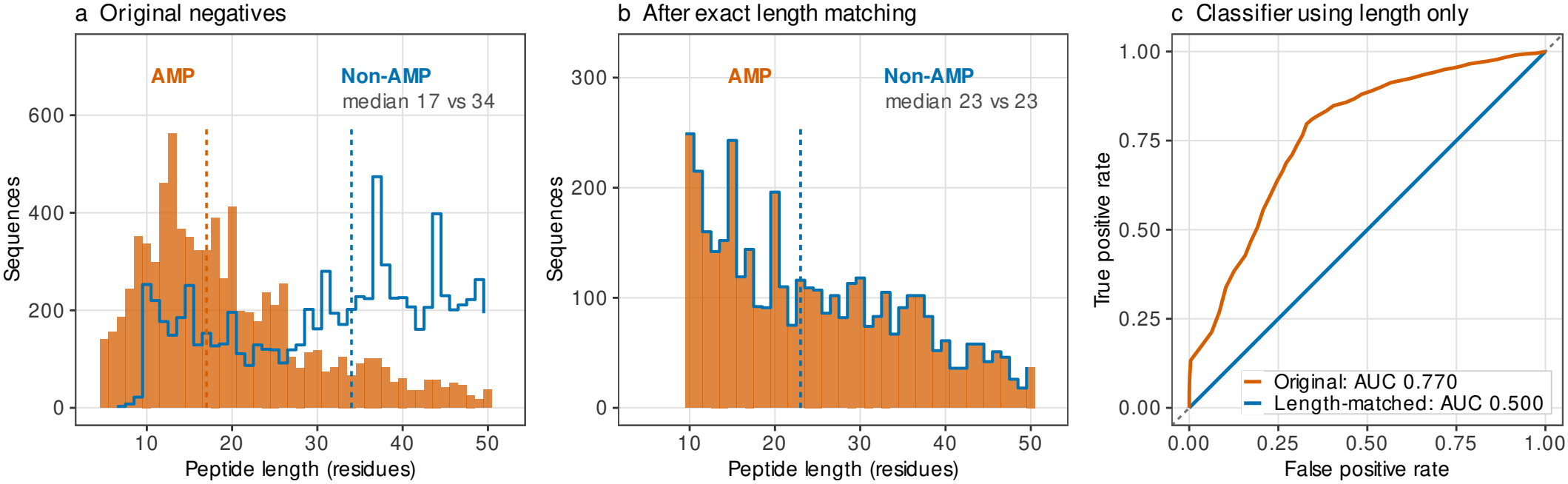
Peptide length as a confound. (a) Length distribution of the original benchmark by class (filled bars, AMP; outline, non-AMP) with class medians marked. Both classes are capped at 50 residues, yet the medians differ twofold. (b) The same after exact 1:1 length matching, where the non-AMP outline coincides with the AMP bars exactly, by construction. (c) ROC of a classifier whose only input is the residue count: AUC-ROC 0.770 on the original benchmark and 0.500 after matching.

### C. Methods under test

The instrument is a validation-AUC-weighted soft-voting ensemble of CNN-Attention, XGBoost, LightGBM and SVM-RBF over a 614-dimensional descriptor vector (amino acid composition, dipeptide composition, composition-transition-distribution, and AMP-specific physicochemical features), with seven classical baselines and three neural architectures for comparison. The lineage of these components in AMP prediction is well established: compositional descriptors with discriminant or kernel classifiers [13], [14], two-level multi-label classifiers [13], deep models for short peptides [15], species-specific zero- and few-shot approaches [16], and more recently protein language model transfer [17], [18]. Related work has extended these strategies to adjacent peptide problems, including cross-family transfer from AMPs to anticancer peptides [19] and unified therapeutic-peptide prediction that feeds sequence length in as an explicit input feature [20], a design choice worth revisiting in light of Section IV-K. Our purpose is not to adjudicate between these families but to show that the protocol choice can dominate the differences between them.

## III. Materials and Methods

The guiding constraint throughout is that the models, features, hyperparameters and training code are identical across protocols. Only the data partitioning and the composition of the negative class change.

### A. The benchmark under test

Sequences were collected from DBAASP, APD6, DRAMP (General and Natural) and UniProt, giving 48,611 raw entries, reduced to 31,266 after cross-database exact and near-duplicate removal, and balanced to 8,084 sequences per class for a final benchmark of 16,168. The positive class comprises DBAASP (6,400), DRAMP General (1,283), APD6 (305), UniProt AMP entries (60) and DRAMP Natural (36); all 8,084 negatives are UniProt entries without an antimicrobial annotation. No sequence carries both labels, but clustering at 40% identity places 312 negatives (3.9%) and 199 positives (2.5%) in the same cluster as a member of the opposite class, across 45 of the 8,714 clusters, which bounds the label noise the UniProt negative-sampling convention introduces. Each sequence is encoded as the 614-dimensional descriptor vector described above, standardised with statistics fit on training data only.

#### The ensemble

The instrument under test is a soft-voting ensemble of four learners over the 614-dimensional descriptor vector: a convolutional network with attention (322,595 parameters), XGBoost, LightGBM and an RBF-kernel SVM with probability estimation enabled. The convolutional component treats the descriptor vector as a single-channel signal of length 614 and applies three convolutional blocks of width 3 (1 to 64 to 128 to 256 channels), each with batch normalisation, ReLU and dropout 0.3, with max-pooling of stride 2 after the first two; squeeze-and-excitation gating then reweights the 256 channels through a bottleneck of 32, and additive attention over positions is concatenated with global average pooling before a fully connected head of 512, 256, 128 and 2 units. It is therefore a nonlinear learner over compositional descriptors rather than a sequence model, since the position axis runs over feature indices. The three classical learners are trained on the training partition only and the network with early stopping on validation loss. Each learner’s predicted probability is weighted in proportion to its validation AUC-ROC and the weighted sum is thresholded at 0.5; Section IV-E shows that this weighting is numerically inert. Full hyperparameters and the trained checkpoints are in the release.

#### Original protocol

A random 70/15/15 split (11,317 / 2,425 / 2,426) at seed 42, with ensemble weights taken from validation AUC and the test partition evaluated once. This is the protocol that produced MCC 0.8338.

### B. Control 1: homology

All sequences were clustered with MMseqs2 [21] at minimum sequence identity thresholds of 20%, 40%, 60%, 80% and 90% (coverage 0.8, `--cov-mode 0`), and train/validation/test partitions were formed over whole clusters, so that no cluster is split across partitions. On the length-matched benchmark, the 8,098 sequences fall into 4,808 clusters at 40% identity, 5,114 at 60%, 5,891 at 80% and 6,723 at 90%; the 40% threshold is therefore the strictest control and the 90% threshold approximates exact-duplicate removal only. Partition sizes are held at the original 70/15/15 proportions. We report 40% as the headline control and the full threshold sweep as the degradation curve.

### C. Control 2: length

We built a length-matched benchmark by pairing each retained positive with a negative of *identical* length, restricted to the 10–50 residue range where both classes have support. This yields 4,049 sequences per class (8,098 total) whose per-class length histograms are exactly equal: 41 distinct lengths, median 23 and mean 24.56 residues in both classes, identical minima and maxima. Matching is exact rather than caliper-based (the bin width is one residue) and is done without replacement, so no negative is used twice and no residual length imbalance remains to be adjusted for. The negative pool is shallow at short lengths, so a positive whose length is already exhausted is dropped rather than matched approximately: 4,049 of the 7,005 positives in the 10–50 range are retained. We note one defect in the shipped benchmark that this exposes: 142 of its 4,049 matched negatives (3.5%) carry UniProt accessions that appear nowhere in the 16,168-sequence source, so the matched benchmark is not strictly a subset of the original one. The redrawn benchmarks of Section IV-J are drawn entirely from the source pool and do not share this defect.

### D. Control 3: sample size

Length matching reduces the benchmark from 16,168 to 8,098 sequences, which confounds the control with a reduction in training data. We therefore constructed a *size control*: the original, unmatched benchmark subsampled at random (seed 42) to exactly the matched benchmark’s per-split, per-class sizes, reusing the same clustering so that subsets of cluster-disjoint partitions remain cluster-disjoint. Any drop observed in the size control is attributable to data volume; any additional drop in the matched benchmark is attributable to length.

This gives three benchmarks: `original` (homology control only), `size_control` (homology plus sample size) and `matched` (homology, sample size and length), crossed with four identity thresholds.

### E. The length-only reference classifier

As a negative control for the benchmark rather than the model, we evaluate a classifier whose only input is the number of residues, fit and evaluated on the same partitions as every other model. Its score bounds the share of measured performance that requires no sequence information at all.

### F. Seeds, cross-validation and prospective evaluation

The main sweep retrains every model under ten seeds (42, 1–9) on each of the three benchmarks at each of four thresholds. The cluster-disjoint partitions are fixed per benchmark and threshold, so seed variation captures training stochasticity (neural initialisation, subsampling) and not partition variance; deterministic learners therefore show zero seed variance, and we report partition variance separately through cross-validation.

#### Cluster-disjoint cross-validation

Clusters are partitioned into five folds by a greedy largest-cluster-first assignment; for fold *k*, fold *k* is the test set, fold *k* + 1 the validation set used for ensemble weighting, and the remainder training. No cluster is split across folds in any fold.

#### Deposition-order split

No stored record carries a curation date, so we use the DBAASP accession number as a proxy for deposition order: of 2,352 positives carrying a parseable DBAASP identifier (range 12–25,320), the oldest form the training set and the most recent 20% the test set, with undated positives assigned to training. This approximates the prospective setting in which a model trained today is applied to peptides characterised tomorrow. We run it with and without additional homology control.

### G. Redrawn partitions

To separate the controls’ costs from the particular draw they were measured on, we repeat the whole decomposition ten times. Each draw independently redraws the length-matched negative set (exact matching, without replacement, from the benchmark’s own UniProt pool), re-clusters that benchmark with MMseqs2 at 40% identity, and redraws the cluster-to-partition assignment for all three benchmarks. Clusters are placed so that each partition meets its 70/15/15 size quota *and* its per-class quota; a size-only rule lets the class ratio drift in opposite directions between the matched and original arms, which biases the between-arm comparison the study depends on.

### H. Stronger sequence baselines

To test whether the descriptor representation, rather than the ensemble, is the limiting factor under control, we add two models that consume sequence directly: (i) **ESM-2 650M** [17] embeddings (mean-pooled) with either a linear probe or Light-GBM on top, and (ii) a **residue-level CNN** trained on one-hot residue sequences with early stopping on validation AUC. Both are evaluated on the identical partitions. The ESM-2 pipeline has no stochastic component once the embeddings are cached and the partition is fixed: neither the linear probe nor the gradient-boosted head subsamples rows or features. We confirmed this by rerunning it under nine further seeds, which reproduce every value in Table IV to six decimal places. The residue-level CNN is stochastic and is reported at seed 42.

### I. Checking the homology control against an exact aligner

MMseqs2 prefilters on shared *k*-mers, which is the regime where short sequences are weakest, and the median peptide here is 23 residues. If the clustering missed homology, every controlled number in this paper would be optimistic. We there-fore re-tested it with exact Smith-Waterman (BLOSUM62, gap open − 11, extend −1) at the same thresholds the clustering used, *≥* 40% identity over *≥* 80% coverage of both sequences: all-against-all on a random 2,000 sequences, and across every test *×* train pair of the matched 40% split.

### J. External benchmark audit

We applied the same two controls to three published, distributed AMP benchmarks (AMP Scanner v2 [22], AMPlify in its balanced and imbalanced forms [23]) by clustering each benchmark’s own train and test partitions with MMseqs2 at 40% identity and (a) removing test sequences sharing a cluster with training data, (b) length-matching the test negatives to the test positives, and (c) both. Because the controls shrink the evaluation set, we subsample to a common per-class size per benchmark and report the mean and standard deviation over repeats. We separately report, for each benchmark’s *own* published split, the fraction of test sequences sharing a 40%-identity cluster with that benchmark’s training data, and the AUC-ROC obtainable from length alone.

### K. Statistics

MCC [24] is the primary metric, being insensitive to the class-balance changes the controls induce. Confidence intervals are 2,000-resample bootstraps of the test partition (seed 42). Paired model comparisons use the exact two-sided binomial (McNemar) test on discordant test predictions. Calibration is reported as Brier score and expected calibration error over ten equal-width bins. Deployment behaviour is reported as precision and recall at AMP prevalences of 50%, 10% and 1%, obtained by reweighting the negative class at a fixed 0.5 threshold. Ensemble subsets are evaluated by enumerating all fifteen non-empty combinations of the four components under both uniform and validation-AUC weighting.

## IV. Results

### A. The original benchmark is substantially separable by length alone

On the uncontrolled 16,168-sequence benchmark, the median AMP is 17 residues and the median non-AMP is 34. A classifier given only the residue count achieves **AUC-ROC 0.7699** (Fig. 2). Under the 40% cluster-disjoint split with the original negatives it achieves **MCC 0.4336**; after exact length matching it achieves **MCC** − 0.0711, i.e. slightly worse than chance, confirming that the matching removes the signal it was designed to remove rather than merely attenuating it.

For context, this length-only classifier would have ranked above several published AMP predictors on reported accuracy. Its score is not a property of any model; it is a property of the benchmark.

### B. The cost of each control, decomposed

Before decomposing that cost, it is worth quantifying what the original protocol leaves open. Under random 70/15/15 splitting of this benchmark, 52.3 *±* 0.9**% of test sequences share a 40%-identity cluster with the training partition** (mean *±* SD over 200 random splits; 47.7 *±* 1.3% of test positives and 56.9 *±* 1.2% of test negatives), and 27.7% still do at 90% identity (Fig. 1a). The benchmark holds 8,714 distinct clusters among its 16,168 sequences at 40% identity, so roughly half of it is family-redundant (Fig. 1b). Every controlled result below uses cluster-disjoint partitions, for which this quantity is 0% by construction. Measured on test positives alone, for comparability with the external audit in Section IV-K, our benchmark’s 47.7% sits inside the 44% to 70% range those three published benchmarks show.

Table I decomposes the ensemble’s reported MCC. Homology control at 40% identity costs 0.0673 at this partition, a figure Section IV-J shows does not survive redrawing. Reducing the data to the matched benchmark’s size costs nothing; the size control is in fact 0.0090 *higher*, within seed noise, so the subsequent 0.1150 drop is attributable to length matching and not to data volume. The net effect of the two controls is −**0.1733 MCC**, or a 21% relative reduction.

**TABLE I.** What each control costs, measured at one partition and over ten redrawn ones. Each row adds one control to the row above it. Only the length term reproduces.

|  | Δ MCC |  |
|---|---|---|
| Evaluation | One partition | Ten redraws |
| Random split, uncontrolled | n/a | n/a |
| + homology control at 40% | −0.0673 | −0.0248 ± 0.0170 |
| + sample-size control | +0.0090 | −0.0117 ± 0.0132 |
| + length matching | **−0.1150** | **−0.1142 ± 0.0574** |
| Net effect of the two controls | −0.1733 | −0.1507 ± 0.0604 |

Ensemble MCC at the single partition is 0.8338 uncontrolled, then 0.7665 ± 0.0082, 0.7755 ± 0.0045 and 0.6605 ± 0.0111 as each control is added; the ± there is over ten training seeds at a fixed split. The right-hand column instead redraws the partition and the length-matched negative sample ten times (Section IV-J). The length term reproduces; the homology term does not.

The degradation is monotone in the strictness of the homology control (Fig. 3). On the matched benchmark the ensemble scores 0.6605 at 40% identity, 0.7159 at 60%, 0.7057 at 80% and 0.7433 at 90%; on `original`, 0.7665, 0.7833, 0.8022 and 0.8124. The 90% threshold, which removes little more than exact duplicates, recovers most of the homology component but none of the length component: the two controls are close to independent, and reporting either alone understates the problem.

**Fig. 3.**
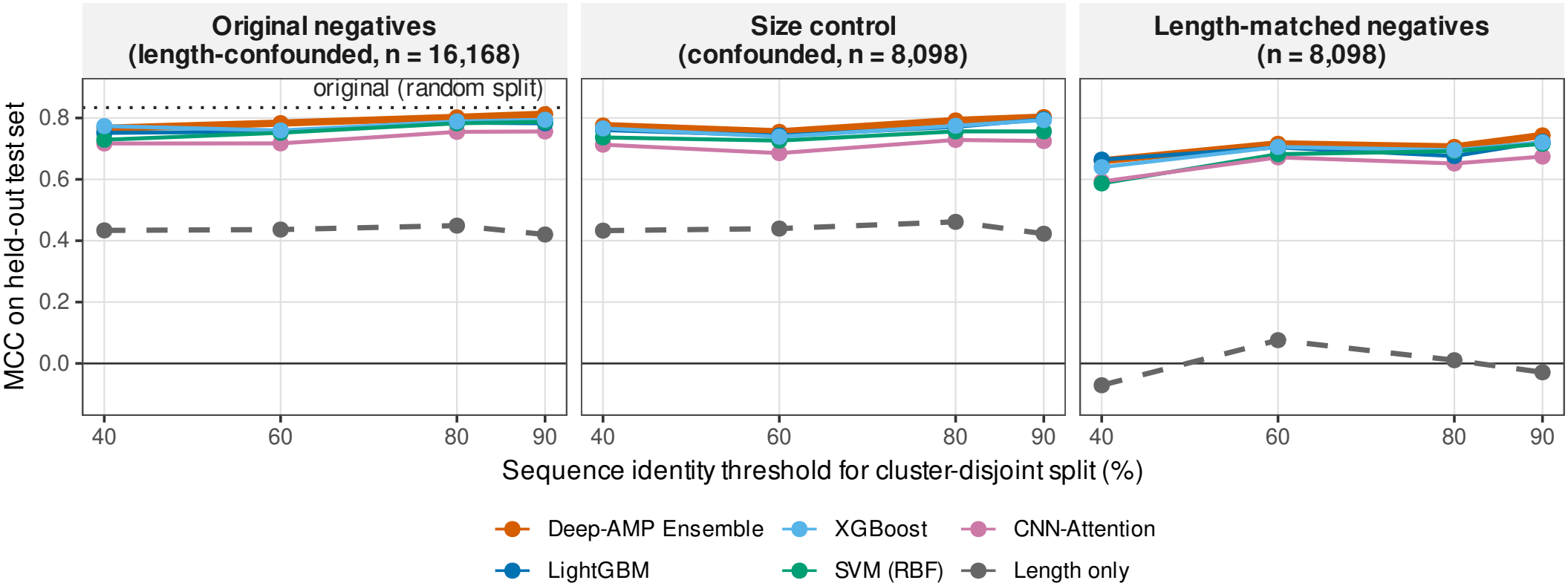
MCC against the clustering identity threshold for the three benchmarks, with the original uncontrolled result marked.

### C. The controls reorder the methods

If the controls acted as a uniform rescaling, the ranking of methods would be preserved and the exercise would be of limited practical consequence. They do not. Table II gives all eleven models plus the length-only reference under the original protocol and under both controls at 40% identity.

The rank correlation between the two columns is Spearman ρ = 0.61 (*p* = 0.047) over all eleven models, and ρ = 0.50 (*p* = 0.14) over the ten baselines alone. The losses range from 0.149 (LightGBM) to 0.262 (Random Forest), a spread of 0.113 MCC, nearly twice the entire spread between the best and worst baseline under the original protocol (0.060).

**TABLE II.** The controls reorder the models. All eleven models and the length-only reference, ranked before and after both controls are imposed.

| Model | Original protocol | Controlled | Δ | Rank, original | Rank, controlled |
|---|---|---|---|---|---|
| Deep-AMP Ensemble | 0.8338 | 0.6605 ± 0.0111 | −0.1733 | 1 | 2 |
| SVM (RBF) | 0.8213 | 0.5868 ± 0.0000 | −0.2345 | 2 | 8 |
| LightGBM | 0.8133 | **0.6644 ± 0.0000** | −0.1489 | 3 | **1** |
| Gradient Boosting | 0.8093 | 0.6417 ± 0.0124 | −0.1676 | 4 | 3 |
| XGBoost | 0.8042 | 0.6397 ± 0.0000 | −0.1645 | 5 | 4 |
| CNN-Attention | 0.7849 | 0.5788 ± 0.0286 | −0.2061 | 6 | 9 |
| CNN-BiLSTM-Attention | 0.7845 | 0.6222 ± 0.0182 | −0.1623 | 7 | 5 |
| CNN-1D | 0.7681 | 0.5568 ± 0.0402 | −0.2113 | 8 | 10 |
| SVM (Linear) | 0.7655 | 0.6148 ± 0.0000 | −0.1507 | 9 | 6 |
| Random Forest | 0.7618 | 0.5000 ± 0.0082 | −0.2618 | 10 | 11 |
| Logistic Regression | 0.7610 | 0.6033 ± 0.0000 | −0.1577 | 11 | 7 |
| *Length only*<sup>†</sup> | *0.4336* | −0.0711 ± 0.0000 | −0.5047 | n/a | n/a |

Original-protocol values are a single run at seed 42, recomputed from the archived per-sequence probabilities at a 0.5 threshold, so they carry no standard deviation. The ensemble's controlled value is the four-model sweep used throughout the paper; the eleven-model sweep that produced the baseline column gives 0.6593 ± 0.0100 for it, a difference of 0.0012 attributable to retraining the stochastic component. Random Forest's controlled mean is 0.49997, not a rounded placeholder. <sup>†</sup>The length-only row is not a random-split value: it is the length classifier under the 40% cluster-disjoint split with the original negatives, against the same split after length matching, so its Δ isolates the effect of the length control on the confound itself.

The most consequential reordering concerns SVM-RBF. Under the original protocol it is the strongest single model and the one the ensemble struggled to beat; under control it loses 0.235 MCC and falls from second to eighth of the eleven models, below linear SVM and logistic regression. Among the ten baselines alone the same change is first to seventh; we quote the eleven-model ranking of Table II throughout. Its apparent strength was disproportionately dependent on the confounded benchmark. Conversely LightGBM, third under the original protocol, is the most robust model tested and becomes the single best model under control, outperforming the ensemble built on top of it. The direction of this result matches García-Jacas *et al*. [26], who found that deep models did not outperform shallow ones for AMP classification under fair comparison; our contribution is to show that the protocol alone reorders them, with the models and features held fixed.

### D. The ensemble’s advantage does not survive control

Under the original protocol the ensemble leads SVM-RBF by 0.0125 MCC. A 2,000-resample bootstrap of that margin gives a 95% interval of [− **0.0071**, +**0.0313**] with *P*(margin *>* 0) = 0.907: the reported improvement was never statistically distinguishable from zero, a point obscured by bootstrapping the ensemble’s MCC rather than the margin.

Under control the margin changes sign. Table III reports the ensemble’s margin over the best single model in each setting, over ten seeds. Fig. 4 gives the seed-level distribution of the margin over LightGBM, which is the best single model on the matched benchmark at every seed, though not on `original`, where XGBoost is.

**TABLE III.** The ensemble’s advantage over its best single component disappears under both controls. Negative values mean the component wins.

| Benchmark | 40% | 60% | 80% | 90% |
|---|---|---|---|---|
| matched | −**0.0039** | +0.0102 | +0.0104 | +0.0207 |
|  | ±0.0111 | ±0.0042 | ±0.0029 | ±0.0046 |
| original | −**0.0070** | +0.0238 | +0.0134 | +0.0182 |
|  | ±0.0082 | ±0.0026 | ±0.0050 | ±0.0046 |
| size_control | +0.0093 | +0.0121 | +0.0175 | +0.0070 |
|  | ±0.0045 | ±0.0046 | ±0.0034 | ±0.0068 |

**Fig. 4.**
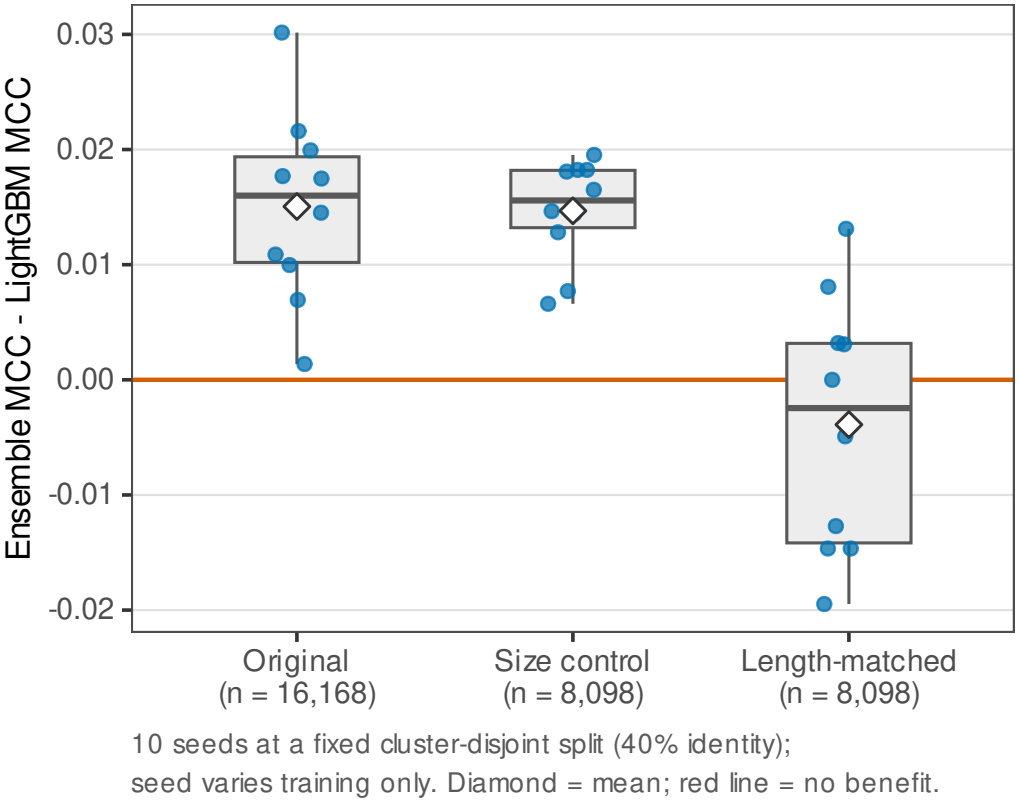
Distribution over ten seeds of the ensemble’s margin over LightGBM at the 40% identity threshold, for each of the three benchmarks; the line at zero marks no benefit and the diamond is the mean. On the length-matched benchmark the box straddles zero.

At the strictest homology threshold the ensemble is behind its best component on both the matched and the original bench-mark; the margin on `matched` at 40% ranges from − 0.0195 to +0.0131 across seeds, straddling zero. The best single model is LightGBM or XGBoost in every controlled setting, never SVM-RBF and never a neural model. Exact McNemar tests against LightGBM on paired predictions confirm that no controlled setting reaches significance: *p* = 0.389 (`matched`, 40%; 29 vs. 37 discordant), *p* = 0.053 (`matched`, 90%; 26 vs. 13), *p* = 0.182 (`original`, 40%; 47 vs. 34) and *p* = 0.099 (`size_control`, 40%; 24 vs. 13).

Five-fold cluster-disjoint cross-validation of the full ensemble, which captures partition variance that the fixed-split sweep cannot, agrees: at 40% identity on `matched`, the ensemble averages 0.6769 *±* 0.0391 against XGBoost at 0.6556 *±* 0.0462 and LightGBM at 0.6531 *±* 0.0334, with fold-to-fold standard deviations roughly twice the margin. At 90% identity the ordering is the same with tighter folds (ensemble 0.7356 *±* 0.0136, XGBoost 0.7221 *±* 0.0158).

### E. The ensemble is not the best configuration of its own parts, and its weighting is inert

Enumerating all fifteen non-empty subsets of the four components on the original protocol, **five outperform the full four-model ensemble** (0.8338): XGB+SVM at 0.8459, LGBM+SVM at 0.8427, CNN+XGB+SVM at 0.8410, XGB+LGBM+SVM at 0.8378 and CNN+LGBM+SVM at 0.8377 (Fig. 5). The best configuration is a two-model pair, and it beats the full ensemble by 0.0122, approximately the same size as the ensemble’s claimed advantage over the best baseline. A selection procedure that chose the best subset on the test partition would have reported a larger “gain” with no modelling change whatsoever.

**Fig. 5.**
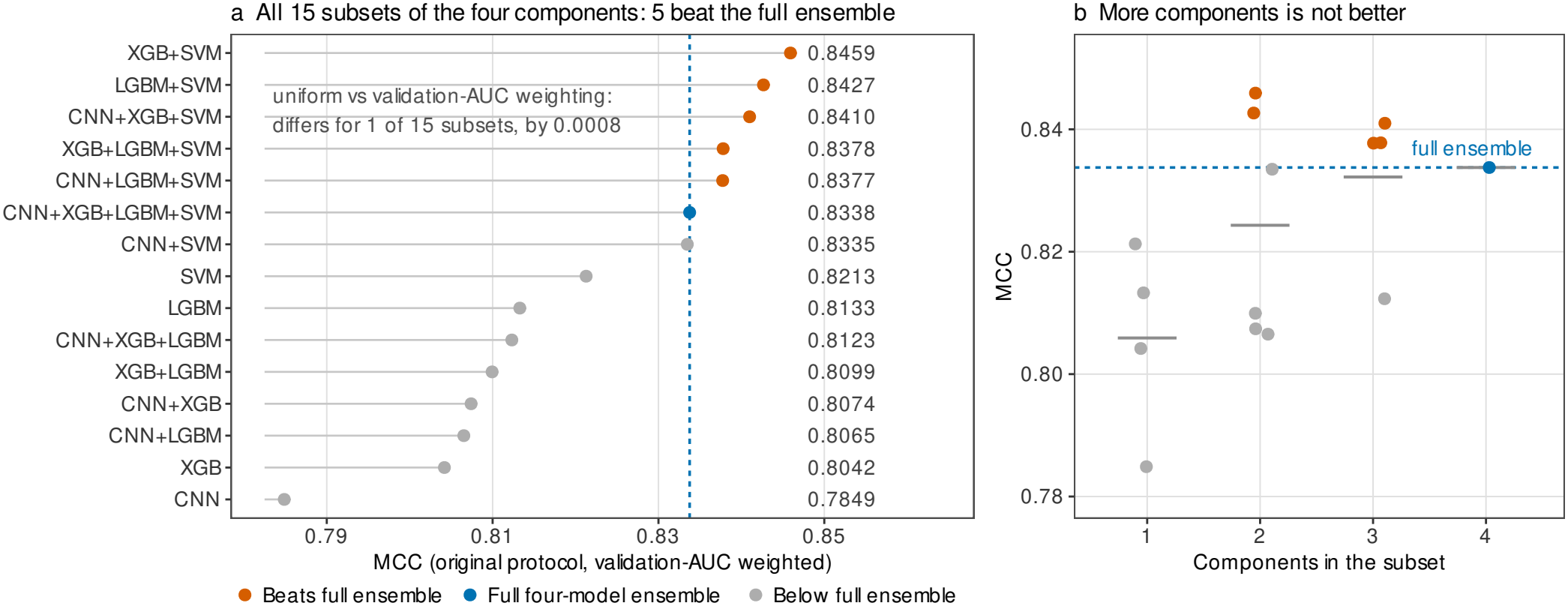
The ensemble is not the best configuration of its own components. (a) All fifteen non-empty subsets of the four base learners, ranked by MCC under the original protocol; the full four-model ensemble is marked and the five subsets that exceed it are highlighted. The inset note reports how far validation-AUC weighting departs from uniform averaging across the fifteen subsets. (b) The same values against the number of components, with per-count means; the best configuration is a two-model pair and the four-model ensemble is exceeded by subsets of every smaller size.

The validation-AUC weighting that the ensemble’s design rests on is numerically inert. The four validation AUCs produce weights of 0.249, 0.250, 0.250 and 0.250, and uniform and validation-AUC weighting give identical MCC for four-teen of the fifteen subsets (the exception, CNN+XGB+LGBM, differs by 0.0008). The “validation-AUC-weighted soft voting” is arithmetic averaging.

### F. Under control, simpler sequence models beat the descriptor ensemble

The uncontrolled protocol supports the conclusion that engineered descriptors with an ensemble are competitive with learned representations. Control reverses it. Table IV compares the descriptor ensemble with two models that consume sequence directly.

**TABLE 4.** Under control, two plainer sequence models beat the descriptor ensemble.

| Benchmark, threshold | Descriptor ensemble | ESM-2 650M + LightGBM | Residue CNN |
|---|---|---|---|
| matched, 40% | 0.6605 | **0.7300** | 0.6971 |
| matched, 60% | 0.7159 | **0.7389** | 0.7192 |
| matched, 80% | 0.7057 | **0.7185** | 0.7304 |
| matched, 90% | 0.7433 | 0.7441 | **0.7552** |
| original, 40% | 0.7665 | **0.8044** | 0.7621 |
| original, 90% | 0.8124 | **0.8236** | 0.8171 |
| size_control, 40% | 0.7755 | **0.7856** | 0.7219 |

The advantage of the ESM-2 pipeline is largest exactly where the controls bite hardest: +0.070 MCC at the strictest setting, shrinking to +0.001 at 90% identity on the matched benchmark. A mean-pooled frozen embedding with a gradient-boosted head, requiring no architecture design, outperforms the four-model ensemble in all seven controlled settings. The ranking of representation strategies is therefore itself protocol-dependent, which is a caution against cross-paper comparison of AMP predictors evaluated under different protocols.

### G. The homology control holds under an exact aligner

Exact Smith-Waterman confirms 1,664 of the 2,010 same-cluster pairs in the sample (82.8%) and flags 1,154 of the 1,996,990 pairs MMseqs2 separates (0.058%). Applied across the cluster-disjoint split itself, 657 of the 1,215 test sequences (54.1%) have at least one training sequence meeting the criterion. That figure looks alarming and is not: at a per-pair rate of 0.058% against 5,668 training sequences, 96.2% of test sequences would meet it by chance, so the split leaves cross-split similarity at 0.56 times the level random pairing of peptides this short produces. The lesson is about the criterion rather than the clustering. A 40% identity threshold over 80% coverage is weak for 23-residue peptides (it asks for roughly eight matched residues), so pairwise filtering saturates and cannot certify a split at these lengths, whereas cluster-level control can and does.

### H. Prospective (deposition-order) evaluation

Under the deposition-order split, with the oldest DBAASP entries as training and the most recent 20% as test (6,218 training and 940 test sequences), the ensemble reaches MCC 0.7833, with SVM-RBF at 0.7750, XGBoost at 0.7752, Light-GBM at 0.7668 and CNN-Attention at 0.6915. Adding homology control (762 test sequences) lowers all models by roughly 0.02: ensemble 0.7639, SVM-RBF 0.7600, LightGBM 0.7585, XGBoost 0.7533, CNN-Attention 0.6866.

Two observations follow. First, the prospective split is less punishing than full length matching, because its negatives retain the original length distribution. Second, the length-only classifier *inverts* under this split, scoring MCC − 0.1953 without homology control and − 0.2329 with it: peptides characterised more recently are longer, so a length heuristic tuned on older data is actively misleading on newer data. A model that has partially learned length as a proxy will degrade in deployment in a way that no random split can reveal.

### I. Calibration and deployment operating points

The ensemble’s probability estimates are better than its components’ by Brier score in three of four controlled settings (e.g. 0.0838 vs. 0.0894 for LightGBM on `original` at 40%), but *worse* by expected calibration error in all four (0.0422 vs. 0.0254 on `matched` at 40%; 0.0519 vs. 0.0189 on `matched` at 90%). Averaging four learners sharpens the ranking slightly while degrading the reliability of the probabilities themselves, so the ensemble cannot be defended on calibration grounds either.

The operating-point analysis is the practical bottom line. At the balanced prevalence the benchmark presents, the controlled ensemble has precision 0.815 at recall 0.843. Reweighting the negatives to an AMP prevalence of 10% gives precision 0.328; at 1%, a plausible rate for untargeted screening, precision falls to **0.0425**. Roughly 24 of every 25 flagged candidates would be false positives. LightGBM alone gives 0.0461 at the same prevalence.

### J. Stability of the decomposition under redrawn partitions

The decomposition of Table I rests on one partition and one length-matched negative sample. Both are draws, and Table III and Fig. 4 already show that split-to-split variation exceeds seed-to-seed variation here. We therefore repeated the whole decomposition ten times, redrawing the cluster-to-split assignment and the length-matched negative set independently on each draw, re-clustering each redrawn benchmark from scratch, and retraining throughout.

The two terms behave differently. **The length term is reproduced**: − 0.1142 *±* 0.0574 over ten draws, against − 0.1150 at the shipped partition. **The homology term is not**: − 0.0248 *±* 0.0170, against − 0.0673 at the shipped partition, which lies outside the range of all ten redraws (− 0.0573 to − 0.0028). The sample-size term straddles zero (− 0.0117 *±* 0.0132), confirming what it was built to test. The net cost of the two controls, −0.1507 *±* 0.0604, is consistent with the −0.1733 of Table I.

Two conclusions follow, and the second qualifies this paper’s own headline. First, the length effect is a property of the benchmark’s construction rather than of any particular split: it survives redrawing both the partition and the negative sample. Second, the homology cost measured at a single partition overstates what redrawing gives, here by a factor of about 2.5, so a decomposition reported from one split should be read as a point on a distribution rather than as the cost of the control. The length-matched arm is also markedly more variable across draws (SD 0.0604 in MCC) than the original arm (0.0170), because length matching roughly halves the benchmark; a study that matches on length should report partition variance rather than seed variance.

### K. The pattern generalises to published benchmarks

The results above could reflect idiosyncrasies of our bench-mark. Table V applies the same two controls to three published, distributed benchmarks, and Fig. 6 shows the result together with the audit of each benchmark’s own split.

**TABLE V.** The same pattern in three published benchmarks. Each is re-evaluated under the controls of Section IV-K.

| Benchmark | As published | Homology-controlled | Length-controlled | Both | Δ (both) |
|---|---|---|---|---|---|
| AMP Scanner v2 [22] (*n* = 383/class) | 0.7973 ± 0.0132 | 0.7389 ± 0.0139 | 0.8022 ± 0.0142 | 0.7444 | −0.0530 |
| AMPlify, balanced [23] (*n* = 221/class) | 0.8226 ± 0.0272 | 0.6163 ± 0.0159 | 0.8283 ± 0.0216 | 0.6432 | −0.1794 |
| AMPlify, imbalanced [23] (*n* = 277/class) | 0.8494 ± 0.0128 | 0.6871 ± 0.0084 | 0.7775 ± 0.0175 | 0.6150 | −0.2344 |

**Fig. 6.**
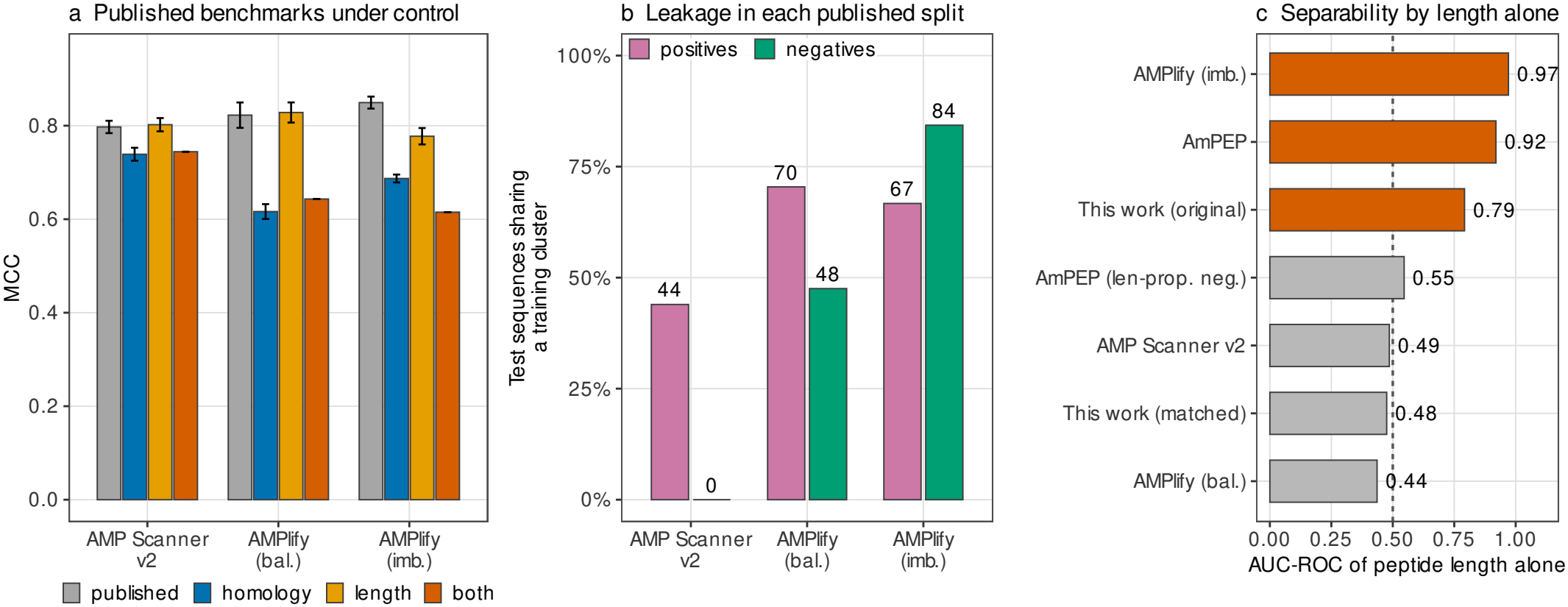
The same controls applied to three published benchmarks. (a) MCC as published and under each control (mean *±* SD over subsampled repeats at a common per-class size), corresponding to Table V. (b) Percentage of each benchmark’s own distributed test partition sharing a 40%-identity cluster with its own training partition, by class. (c) AUC-ROC obtainable from peptide length alone on each benchmark’s own split, ordered; bars above 0.6 are highlighted and the dashed line marks chance.

Homology control costs 0.058 to 0.206 MCC on these benchmarks. Length control costs essentially nothing on the two balanced ones and 0.072 on the imbalanced one, which is consistent with their construction, and is the point of Table VI: these benchmarks differ in *which* confound they leave open, and auditing only one of the two would clear two of the three.

**TABLE VI.** Which confound each benchmark leaves open. Train/test cluster sharing and length separability, measured on each benchmark’s own distributed split.

| Benchmark, split as distributed | Test positives sharing<br>a training cluster | Test negatives sharing<br>a training cluster | Median length<br>(pos / neg) | AUC of<br>length alone |
|---|---|---|---|---|
| AMP Scanner v2 [22] | 313/712 (**44.0%**) | 0/712 (0%) | 28 / 28 | 0.486 |
| AMPlify, balanced [23] | 586/832 (**70.4%**) | 396/833 (47.5%) | 25 / 27 | 0.436 |
| AMPlify, imbalanced [23] | 555/832 (**66.7%**) | 21,657/25,683 (84.3%) | 25 / 131 | **0.971** |
| AmPEP [25], random 70/30 | n/a | n/a | 32 / 160 | **0.920** |
| AmPEP, length-proportional negatives | n/a | n/a | 32 / 37 | 0.546 |
| This benchmark, random 70/15/15 | 47.7 ± 1.3% | 56.9 ± 1.2% | 17 / 34 | 0.770 |
| This benchmark, cluster-disjoint 40% | 0% by construction | 0% by construction | 17 / 34 | 0.770 |
| This benchmark, length-matched | 0% by construction | 0% by construction | 23 / 23 | 0.500 |

Length separability is a property of a benchmark’s class composition rather than of a particular partition, so the last column is computed on each benchmark in full. AmPEP distributes a corpus and a sampling protocol rather than a fixed train/test partition, so its cluster-sharing columns are not defined.

Between 44% and 70% of the test positives in these distributed splits share a 40%-identity cluster with their own training data, which is the mechanism behind the homology column of Table V. Our own benchmark under random splitting sits inside that range at 47.7% of test positives, so this is a shared property of the field’s benchmarks rather than a defect peculiar to any one of them. The length column separates cleanly: AMP Scanner v2 and AMPlify-balanced are effectively length-controlled already (length AUC 0.49 and 0.44) and lose nothing to that control, whereas AMPlify-imbalanced and AmPEP’s random split are severely confounded (0.97 and 0.92). AmPEP appears in the length audit of Table VI but not in Table V: it distributes a single imbalanced corpus rather than a fixed train/test partition, so there is no published split for us to re-cluster, only a sampling protocol to re-run. It is worth including even so, because its own length-proportional negative sampling reduces its length AUC from 0.920 to 0.546, the one example in this audit of the fix being applied in practice, and evidence that it works.

## V. Discussion

### A. What the controls cost, and why it matters

The headline finding is not that our ensemble is bad. Under full control it still reaches MCC 0.66, far above the length baseline of − 0.07, so the models are learning genuine sequence signal. The findings that matter are structural.

First, the magnitude: 0.17 MCC, or 21% relative, is larger than the entire reported difference between the best and worst method in most AMP comparison tables, including our own. Any cross-paper comparison between methods evaluated under different protocols is therefore uninterpretable, because the protocol term dominates the method term.

Second, the direction-changing: because the controls cost different models different amounts (0.149 to 0.262 MCC), they reorder methods. A method selected as best on an uncontrolled benchmark, SVM-RBF here, can be the worst choice under control. This converts the issue from one of calibration (“all numbers are inflated, subtract a constant”) to one of validity (“the comparison does not identify the better method”).

Third, the self-application: the controls dissolved our own contribution. The ensemble’s advantage over its best component, already not significant under the original protocol (bootstrap margin interval [− 0.0071, +0.0313]), reverses sign at the strictest control, is beaten by five of fifteen subsets of itself, and relies on a weighting scheme that is arithmetically equivalent to averaging. We report this because presenting the same ensemble as a method contribution would require the uncontrolled protocol to be taken at face value, and the measurements above show it cannot be.

### B. A minimum reporting standard

From the measurements above we suggest six items, each cheap relative to training a model:

1. **Report a length-only baseline**. One number, computable in seconds. If it is high, the benchmark rather than the method is doing the discriminating.
2. **Report train/test cluster sharing** at a stated identity threshold for the split actually used, as in Table VI.
3. **Split on identity clusters, not sequences**, and state the threshold. Report the degradation curve across thresholds rather than a single point, since the 90% and 40% thresholds differ by 0.08 MCC here.
4. **Match the negative class on length**, exactly where sample size permits, and report a sample-size control so the resulting drop is attributable.
5. **Bootstrap the margin, not the metric**. A confidence interval on a model’s MCC says nothing about whether it beats another model; our own reported margin interval contained zero.
6. **Report an operating point at a realistic prevalence**. A method with MCC 0.66 on a balanced benchmark has precision 4% at 1% prevalence, and prospective users need that number.

### C. Limitations

Several caveats bound our conclusions. The ten-seed sweep varies training stochasticity but not the partition: the cluster-disjoint splits are fixed per benchmark and threshold, which is why deterministic learners show zero seed variance in Table II. Partition variance is addressed separately by five-fold cluster-disjoint cross-validation, which is available for the matched benchmark at 40% and 90% only, and by the ten redrawn partitions of Section IV-J. The latter shows that the homology term in particular is partition-sensitive, so the single-partition figure in Table I should not be quoted on its own. The ESM-2 comparison in Table IV is deterministic given the fixed partition and cached embeddings, so seed replication can neither strengthen nor weaken it; the uncertainty that matters for those rows is partition variance, which only cross-validation would address and which we have not run for these models. The residue-CNN comparison is genuinely single-seed and stochastic, so its small differences at 80–90% identity should not be over-read. The deposition-order split uses accession number as a proxy for date, which is monotone in deposition but not a calendar date, and covers only the DBAASP-derived positives. Our length matching is exact and therefore discards roughly half the data; we have controlled for the resulting size reduction but not for any change in the composition of sources the matching induces. The negative class remains UniProt entries without an antimicrobial annotation, so un-annotated AMPs may be labelled negative; this affects all protocols equally and is not addressed here. Finally, we evaluate binary AMP/non-AMP discrimination only, not activity magnitude, target specificity or mechanism, and no prediction here has been experimentally validated.

### D. Threats to validity

The audit of published benchmarks (Tables V and VI) repartitions and subsamples data distributed by their authors, and so does not reproduce those papers’ own training procedures; the numbers characterise the benchmarks under our controls, not the published methods’ performance as their authors measured it. Our clustering uses one tool at one coverage setting, and alternative clustering choices would shift the thresholds somewhat, though the qualitative monotonicity should be robust; Section IV-G checks the tool itself against exact alignment. We also note an artifact of the original implementation, documented in the released code: the two SVMs were fitted with probability estimation enabled, so `predict` and a 0.5 threshold on `predict_proba` are not the same classifier and disagree on 29 of 2,426 test sequences. All values in this paper use probability thresholding consistently, which is why our recomputed SVM-RBF MCC is 0.8213 rather than the 0.8219 obtained from `predict`. The discrepancy is smaller than, but of the same order as, the margin the original comparison rested on, itself an argument for recomputation from archived probabilities.

## VI. Conclusion

We treated a four-model AMP ensemble of our own as an instrument under test and measured what two routinely omitted evaluation controls cost it. Cluster-disjoint splitting at 40% sequence identity costs 0.067 MCC at the partition we report; exact length matching of the negative class costs a further 0.115, with sample size ruled out as the cause; together they take a reported MCC of 0.834 to 0.661. Repeating the decomposition over ten redrawn partitions and negative samples reproduces the length term (− 0.114 *±* 0.057) but not the homology term (− 0.025 *±* 0.017), so of the two it is length that is a property of the benchmark rather than of the split. The controls are not a rescaling: they reorder the eleven models tested (Spearman ρ = 0.61), demote the strongest baseline from second to eighth of eleven, dissolve the ensemble’s advantage over its own best component, and reverse the conclusion that engineered descriptors are competitive with learned representations. Auditing three distributed benchmarks reproduces the effect and locates its mechanism in 44–70% train/test cluster sharing. At a realistic 1% screening prevalence, the controlled model’s precision is 4.3%.

The constructive reading is that both controls are cheap, that one of them (length) is almost free to report as a single baseline number, and that the field’s benchmarks differ in which confound they leave open, so an audit is required per benchmark rather than once. We release the benchmark, the clustering, the per-sequence predictions and a verification script so that each number above can be recomputed without retraining.

## Data and Code Availability

The benchmark, MMseqs2 cluster assignments at five identity thresholds, analysis and figure scripts, trained models, persequence predictions of every run, and the controlled sweep (3 benchmarks *×* 4 thresholds *×* 10 seeds) will be released publicly, with a permanent DOI, on acceptance, and are available to reviewers on request during review. A script, `verify.py`, recomputes the original-protocol MCC and AUC, the length-only AUC and median lengths, the subset enumeration, and the three-way decomposition of Table I directly from the archived artifacts; it reports ten checks and exits non-zero on any mismatch. The 141 MB descriptor matrix is omitted and regenerated by the feature-extraction code from the released sequences. Bootstrap intervals recomputed from the archived probabilities agree with those from the original analysis to within 0.003 MCC. The dataset is *derived* from DBAASP, APD6, DRAMP and UniProt; each source carries its own terms, and reusers should consult the original databases.

